# Neotype designation and re-description of Forsskål’s reticulate whipray *Himantura uarnak*

**DOI:** 10.1101/2020.10.22.350629

**Authors:** Philippe Borsa, Collin T. Williams, Ashlie J. McIvor, Michael L. Berumen

## Abstract

A serious impediment to the taxonomy of the reticulate whipray *Himantura* spp. species complex has been the absence of a type specimen for P. Forsskål’s *H. uarnak*. Here, reticulate whipray specimens were sampled from the Jeddah region, the assumed type locality of *H. uarnak*, and characterized genetically at the cytochrome-oxidase subunit 1 (*CO1*) locus. One of these specimens now in the fish collection of the California Academy of Sciences in San Francisco was designated as neotype. A maximum-likelihood phylogeny of all available *CO1* gene sequences from the genus *Himantura* had the following topology: ((*H. leoparda, H. uarnak*), (*H. undulata*, (*Himantura* sp. 2, (*H. australis* + *Himantura* sp. 1))), *H. tutul*), where *H. uarnak* haplotypes formed a distinct sub-clade sister to *H. leoparda*. Based on these *CO1* gene sequences, the geographic distribution of *H. uarnak* includes the eastern Mediterranean, the Red Sea, the East African coast, and the Arabian Sea. Two lineages in the reticulate whipray species complex remain to be named.

**Notice:** The present article in portable document (.pdf) format is a published work in the sense of the International Code of Zoological Nomenclature [International Commission on Zoological Nomenclature (ICZN)1999]. It has been registered in ZooBank (http://zoobank.org/), the online registration system for the ICZN. The ZooBank life science identifier for this publication is urn:lsid:zoobank.org:pub:B2113697-5EBF-4364-B50C-63019A1A076A. The online version of this work is archived and available from the bioRxiv (https://biorxiv.org/) repository.

## Introduction

Among the marine fishes listed from P. Forsskål’s expedition to the Red Sea were three stingrays, namely the cowtail stingray *Pastinachus sephen* (Fabricius, 1775), the reticulate whipray *Himantura uarnak* (Gmelin, 1789) and the blue-spotted ribbontail ray *Taeniura lymma* (Fabricius, 1775). All three species have long been believed to have a wide Indo-Pacific distribution (e.g., Last & Compagno 1999) but this view has recently been challenged by molecular phylogenetics (Naylor et al. 2012; Arlyza et al. 2013). The type locality for these species is assumed to be Jeddah, the place where P. Forsskål has described most of its fish material from the Red Sea (Fricke 2008). *H. uarnak* has long been understood as the whipray species having densely and regularly spaced, solid, round or oblong dark spots all over the surface of the dorsal side (Rüppell 1835; Duméril 1865; Randall 1983, 1995; Last and Compagno 1999; Manjaji 2004; Arlyza et al. 2013; Manjaji-Matsumoto et al. 2016). *H. uarnak* belongs to a species complex now found to comprise at least six distinct lineages also including *Himantura undulata* Bleeker, 1852, *Himantura leoparda* Manjaji-Matsumoto and Last, 2008, *Himantura australis* Last, White and Naylor, 2016, *Himantura tutul* Borsa, Durand, Shen, Arlyza, Solihin and Berrebi, 2013, previously under *H. leoparda*, and one lineage, ‘*uarnak 4*’ that still has to be named formally (Naylor et al. 2012; Last et al. 2016; Borsa 2017). To date, no material from the Red Sea has been examined genetically. Reticulate whiprays have recently been reported from the eastern Mediterranean Sea (Lessepsian migrants) and identified as either *H. uarnak* based on spot patterns (Ali et al. 2010) or *H. leoparda* based on nucleotide sequences (Yucel et al. 2017). In the absence of type material, it is unclear which of these lineages – or eventually another lineage – is P. Forsskål’s *H. uarnak*, emphasizing the need for genetically examining reticulate whipray material from the Red Sea (Naylor et al. 2012). Efforts to stabilize the group’s taxonomy by designating a neotype for *H. uarnak* is indeed a task that authors who previously dealt with the taxonomy of species in the genus *Himantura* have neglected (Manjaji 2004; Manjaji-Matsumoto and Last 2008; Borsa et al. 2013; Last et al. 2016; Borsa 2017). The purpose of the present note is to characterize reticulate whipray material collected recently from the Red Sea and designate a neotype for *H. uarnak* in an attempt to clarify the intricate taxonomy of the reticulate whipray species complex.

## Materials and methods

Thirteen reticulate whipray individuals were captured on shallow-water sand flats of the eastern shore of the central Red Sea between 2016 and 2019. Eight individuals were captured near the coastal town of Thuwal, Saudi Arabia (22.313°N 39.090°E); five individuals were captured approximately 15 km offshore on Sirrayn Island (19.616°N 40.647°E) in the Farasan Banks region of Saudi Arabia (Fig. 1). Specimen identification numbers and sampling details are provided in Supplementary Table S1. All live specimens were photographed, measured, and a small section of the pelvic fin was excised as genetic material. One individual [no. KAUST-RSRC-H006 (H006)] was retained whole as voucher specimen to be deposited in a museum collection; it was euthanized by immersion in ice-refrigerated seawater; so was another specimen (H009) whose jaws and tissue sample were preserved. The other individuals were released at their capture site. An additional tissue sample was obtained from an individual on sale at the Jeddah (Saudi Arabia) fish market. The total length, disc width (DW), and length from snout to origin of cloaca were measured on each live specimen. Additional measurements were made on the voucher specimen using the indications of Manjaji (2004). Pigmentation was described using the following parameters: (i) number of spots counted in a rectangular band drawn between spiracles; (ii) number of spots crossed along a line running from mid-scapular point to extremity of pectoral fin, as shown in figure 1 of Borsa et al. (2013); (iii) dorsal-spot diameter; (iv) thickness of paler outer disc margin at extremity of pectoral fin on the dorsal side; and (v) thickness of pigmented outer disc margin at extremity of pectoral fin on the ventral side. Tissue samples (*N* = 14) were placed in ∼95% ethanol and stored at -20°C until further processing.

**Fig. 1.**
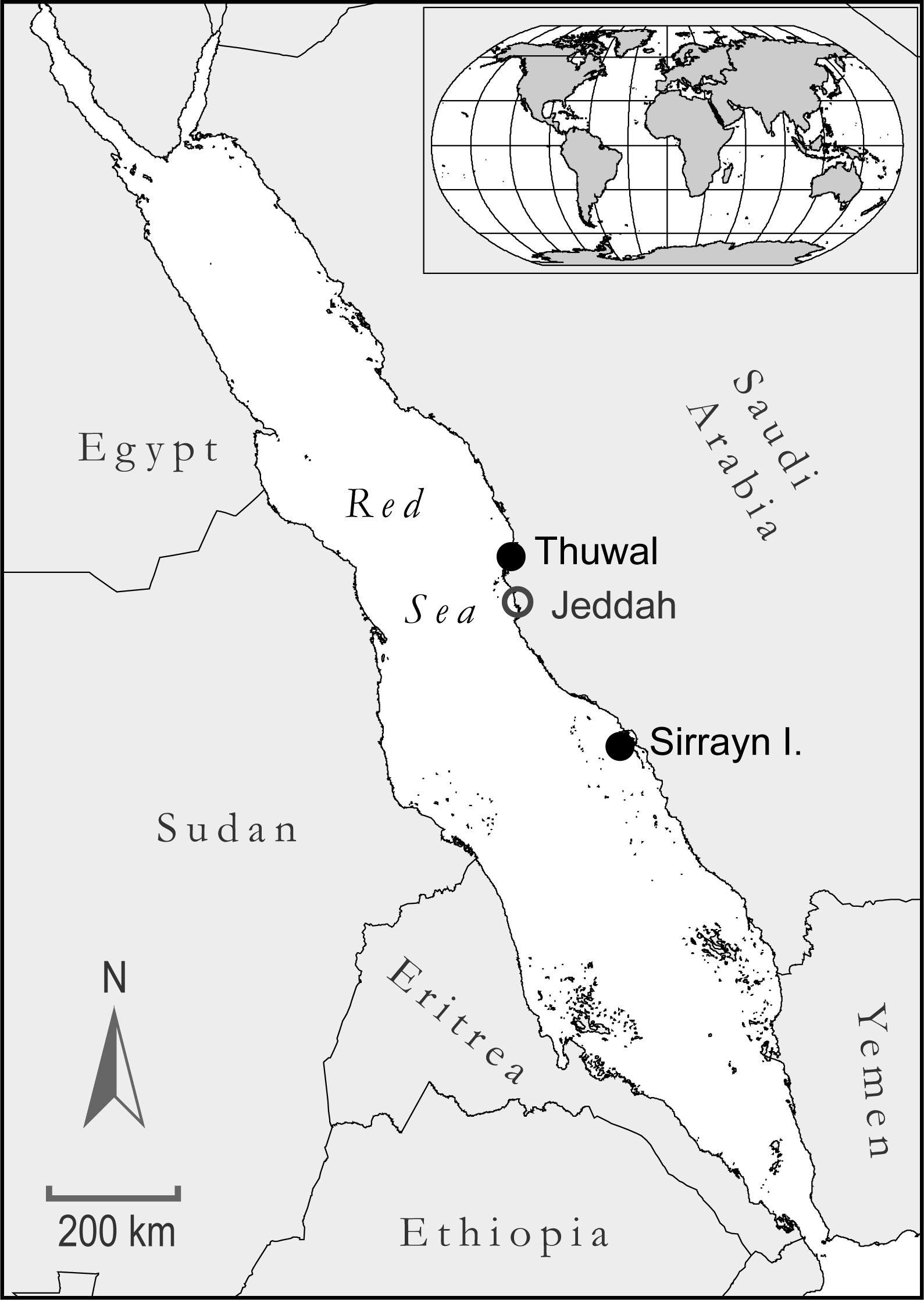
Reticulate whipray sampling localities in the central Red Sea. Jeddah is assumed to be the type locality of Forsskal’s (1775) *H. uarnak* (Fricke 2008)

Genomic DNA was extracted using the DNeasy Blood and Tissue Kit (Qiagen, Valencia, CA) following the manufacturer’s recommendations. The *FishF1*/*FishR1* primer pair (Ward et al. 2005) was used to target a 655-bp portion of the *COI* gene located between homologous nucleotide sites no. 5571 and no. 6225 of the mitochondrial DNA in *H. leoparda* (NC_028325; Shen et al. 2016) for amplification by polymerase chain reaction (PCR). PCR was run in individual wells loaded with 12 μL reaction mixture composed of 2 μL DNA extract, 5.4 μL Multiplex PCR Master Mix (Qiagen, Valencia, USA), 0.4 μL of each primer (10 μM), and 3.7 μL H_2_O. The PCR program included a denaturing step at 95°C for 15 min, followed by 35 cycles of denaturing at 94°C for 30 s, annealing at 50°C for 60 s, and extension at 72°C for 45 s, ending with by a final extension step at 72°C for 10 min. The PCR products were analyzed with the QIAxcel DNA Screening Kit (Qiagen) to confirm successful amplification. Amplified DNA was purified by adding 1.2 µL of ExoProStar (GE Healthcare, Piscataway, USA) to each well before a single thermal cycle of 37°C for 60 min, 85°C for 15 min, and a holdat 4°C until sequencing. PCR samples were then sent for Sanger sequencing at the KAUST Bioscience Core Lab. The sequencing reaction was done in each direction, using either *Fish F1* or *Fish R1* as the sequencing primer. Sanger sequencing of purified target DNA was conducted using the 3730xl DNA Analyzer (Applied Biosystems, Foster City, USA).

Reticulate whipray *CO1* gene sequences from the Red Sea were compared to all homologous *Himantura* spp. sequences available including 201 sequences downloaded from the GenBank (www.ncbi.nlm.nih.gov/genbank/; Clark et al. 2016) and BOLD (http://www.boldsystems.org/; Ratnasingham and Hebert 2007) public repositories, and two unpublished sequences (Supplementary Table S2). Reticulate whipray, *Himantura* spp. *CO1* gene sequences from GenBank and BOLD were labelled “*H. astra*” (*N* = 1), “*H. australis*” (*N* = 1), “*H. fava*” (*N* = 1), “*H. leoparda*” (*N* = 85), “*H. tutul*” (*N* = 30), “*H. uarnak*” (*N* = 62), “*H. uarnak* i” (*N* = 2), “*H. uarnak* ii” (*N* = 3), “*H. uarnak* iii” (*N* = 3), “*H. undulata*” (*N* = 11), “*Himantura* sp.” (*N* = 1) and “*Urogymnus asperrimus*” (*N* = 1) (Supplementary Table S2). Partial nucleotide sequences of the *CO1* gene were aligned and trimmed to a core length of 613 bp using the BioEdit sequence editing software (Hall 1999). The nucleotide sequences were sorted by species and nucleotide synapomorphies were then assessed visually.

A maximum-likelihood tree of partial *CO1* gene sequences was constructed using the MEGA X package (Kumar et al. 2018). All three codon positions were included. Of all models tested under MEGA X, the Hasegawa-Kishino-Yano model (Hasegawa et al. 1985) had the lowest Bayesian information score. The evolutionary history of individual haplotypes was inferred by using the Maximum-Likelihood (ML) method (Felsenstein 1981). A discrete Gamma distribution was used to model evolutionary rate differences among sites (*G* = 1.19). Some sites were allowed to be evolutionarily invariable. GenBank sequences nos. EU398838, EU398867, EU398869, JF493649, KF899470, KF899491, KM073002, KM073006 and KP641389 (*Maculabatis* spp.), EU398849, KF899499, KM072993, KM073009 and KM073010 (*Pateobatis* spp.), EU398860, FJ384709, KF899521, KF899528, KM072995 and KR003772 (*Brevitrygon* spp.) and KF965292 were included as an outgroup to root the tree. Mean nucleotide distances between and within lineages were estimated using the Maximum Composite Likelihood model under MEGA X.

## Results

Disc width ranged between 284 mm and 748 mm. The claspers of the one of the five male specimens (HC256) were calcified. Reticulate whipray specimen no. H006 is presented in Fig. 2a-d. The colour of the dorsal side of all individuals sampled was beige and the ventral side was creamy white. The dorsal side was covered with numerous brown spots, which generally were evenly distributed and had a circular, or slightly elongated, or dumbbell shape. In a proportion of individuals, circular spots were often associated as pairs. Colour tones and spot patterns on the dorsal side were similar to those encountered in the reticulate whipray’s mangrove habitat in the central Red Sea (Fig. 2e). The ventral side was either immaculate or with few dark-grey spots, apart from the outer margin of the posterior half of the disk which had a pale greyish hue with densely distributed small, dark-grey spots. A summary of the measurements used to describe spot patterns on all twelve specimens is reported in Table 1. Dorsal-spot diameter was correlated with disk width (Pearson’s *r* = 0.66; *P* < 0.05) and so was dorsal-spot density estimated from transects (*r* = 0.82, *r* = 0.56; Fisher’s combined *P* < 0.01), indicating that both the size and number of dorsal spots increase with age.

**Table 1.**
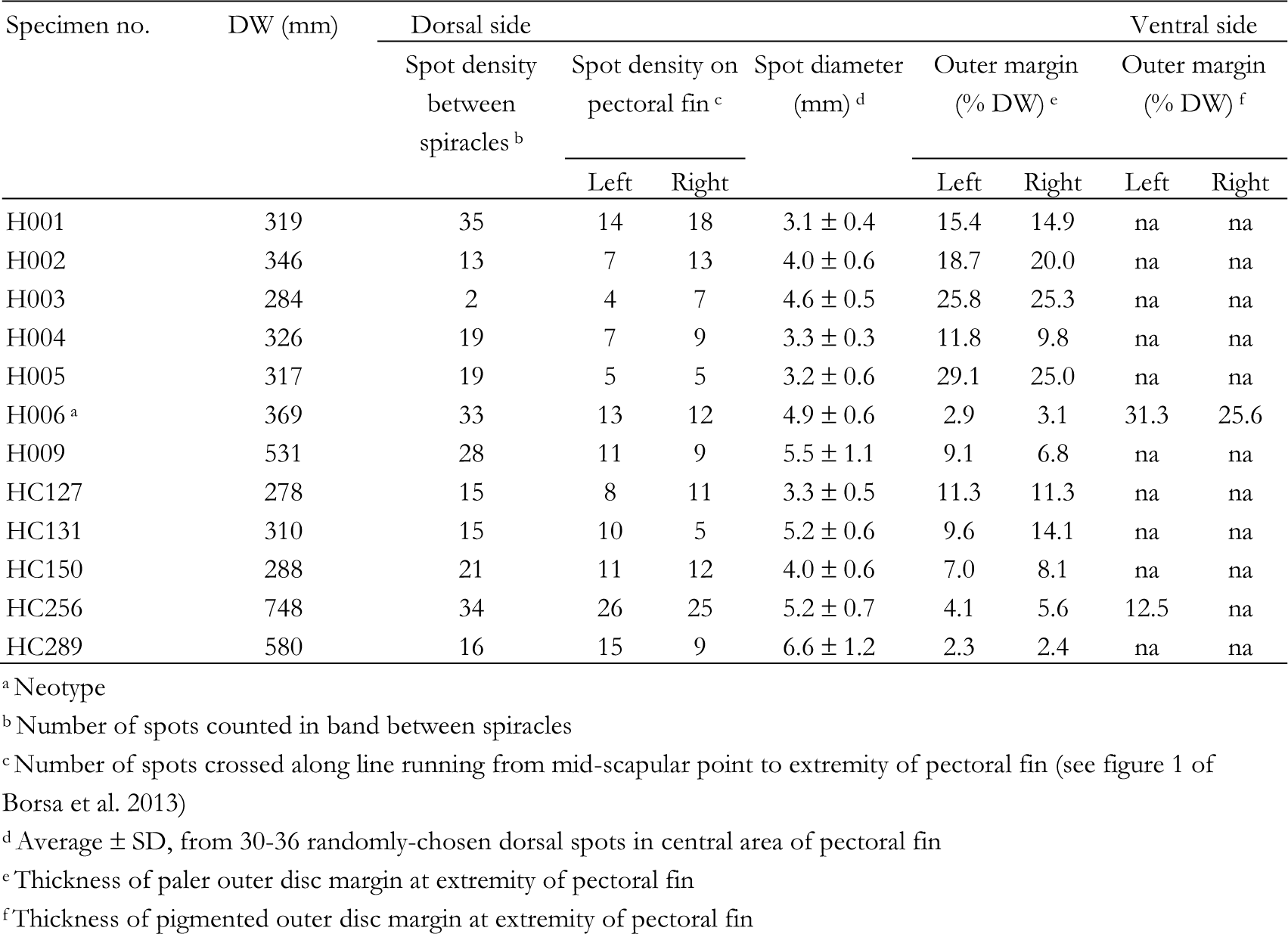
*Himantura uarnak*. Pigmentation patterns in 12 DNA-barcoded specimens sampled from the Jeddah region. *DW* disk width; *na* no data

**Fig. 2.**
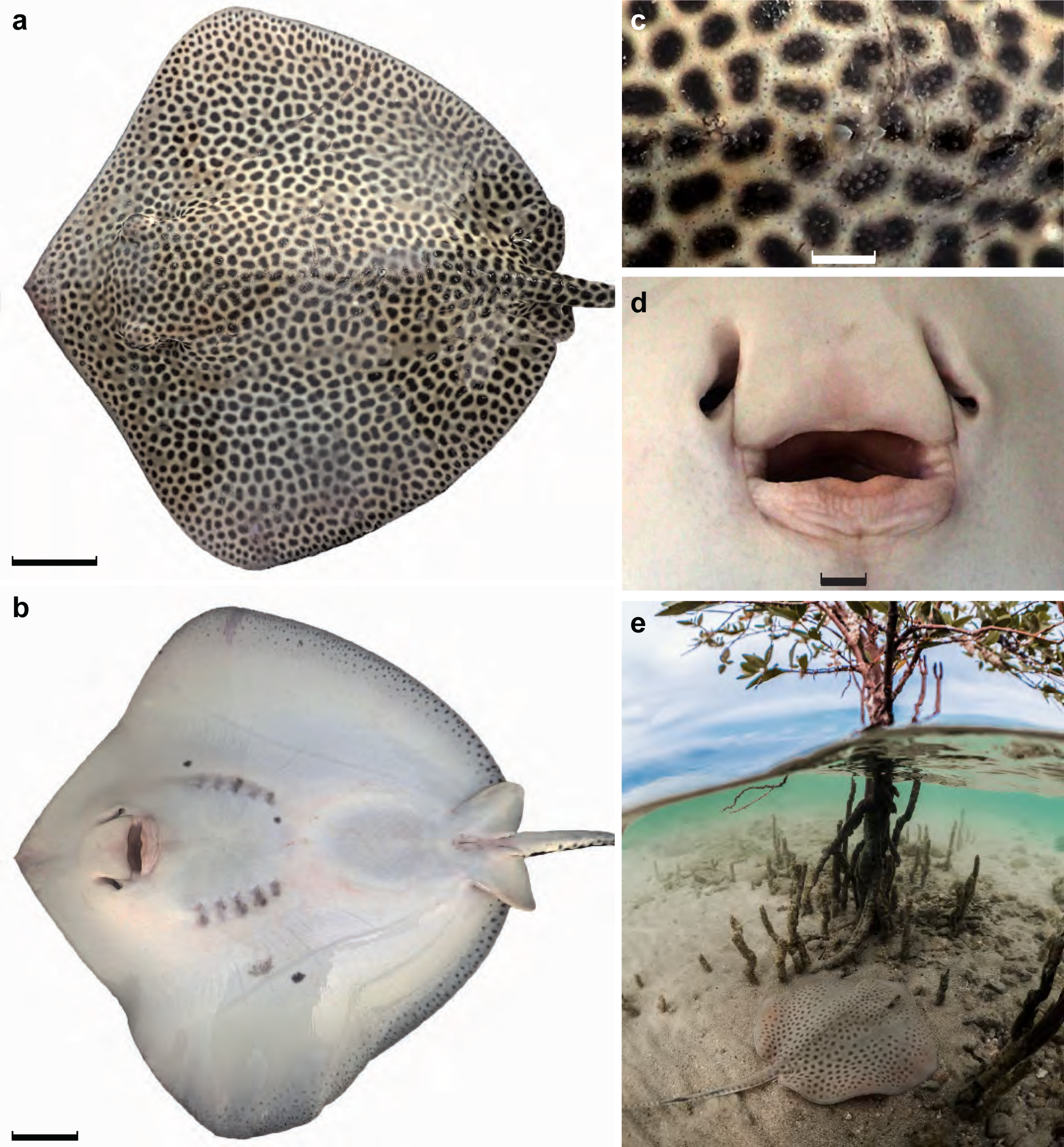
Reticulate whipray *Himantura uarnak* (Gmelin [ex Forsskål], 1789). **a** Dorsal view of neotype, no. CAS-ICH 247241 deposited at the fish collections of the California Academy of Sciences, San Francisco (formerly at the Red Sea Research Center, Thuwal as no. KAUST-RSRC-H006); scale bar: 5 cm. **b** Neotype, ventral view; scale bar: 5 cm. **c** Neotype, spine details in median dorsal region; scale bar: 1cm. **d** Neotype, mouth details; scale bar: 1cm. **e** Sandy mangrove habitat of *H. uarnak* off Thuwal, Red Sea, 22.313°N 39.090°E (credit: M. Bennett-Smith / KAUST)

The ML tree of *Himantura* spp. sequences (Fig. 3) showed four main clades (clades *I-IV*; Arlyza et al. 2013) with strong statistical support. Clade *I* comprised haplotypes sampled from off eastern Africa, including the holotype of *H. tutul*, to the Sulu Sea. Clade-*I* haplotypes were previously identified as either “*H. leoparda*” (Lim et al. 2015; Bineesh et al. 2017; Mohd-Arshaad & Jamaludin 2018), “*H. tutul*” (Arlyza et al. 2013; Borsa et al. 2013), “*H. uarnak*”, “*H. uarnak* i”, “*H. uarnak* ii”, “*H. uarnak* iii” (Bineesh et al. 2017; Gouws 2020; Hanner 2020), “*Himantura* sp.” (Hanner 2020), or “*Urogymnus asperrimus*” (Priyanga et al. 2013). Clade *II* comprised haplotypes sampled from the Arabian Sea to the Sulu Sea, all identified as either “*H. fava*” (Ward et al. 2005) or “*H. undulata*” (Arlyza et al. 2013; Lim et al. 2015; Bineesh et al. 2017; Mohd-Arshaad & Jamaludin 2018; Segura-Garcia & Yain Tun 2018; Ahmed et al. 2019; present work). Three subclades or haplogroups were distinguished within clade *III*. Subclade *III*_*1*_ included two sequences sampled from the Sulu Sea so far labelled “*H. uarnak*” and “*H. uarnak* ii”, respectively (Lim et al. 2015; Hanner 2020) while subclade *III*_*2*_ included haplotypes sampled from tropical Australia exclusively and so far referred to as either “*H. astra*” (Cerutti-Pereyra et al. 2012), “*H. australis*” (Appleyard 2020), “*H. leoparda*” (Cerutti-Pereyra et al. 2012), or “*H. uarnak*” (Cerutti-Pereyra et al. 2012; Appleyard 2020; McGrouther 2020). Haplogroup *III*_*3*_, which in the ML tree was paraphyletic with *III*_*2*_, exclusively comprised haplotypes sampled from the Coral Triangle and labelled either “*H. uarnak*” (Arlyza et al. 2013; Santos et al. 2014; Lim et al. 2015; Mohd Arshaad & Jamaludin 2018; Appleyard 2020; Hanner 2020; Mohd Arshaad 2020) or “*H. uarnak* i” (Hanner 2020). All *Himantura* spp. *CO1* gene sequences sampled from the Red Sea clustered into a distinctive lineage within clade *IV*, hereafter designated subclade *IV*_*2*_, also including haplotypes from the eastern Mediterranean Sea, Natal (South Africa) and the Arabian Sea. These were initially assigned to either “*H. leoparda*” (Bineesh et al. 2017; Yucel et al. 2017) or “*H. uarnak*” (Steinke et al. 2011). Subclade *IV*_*1*_ which was sister to *IV*_*2*_ comprised haplotypes from India, Sunda, Western Australia and Taiwan. These were initially identified as either “*H. leoparda*” (Arlyza et al. 2013; Shen et al. 2016; Appleyard 2017; Ravi et al. 2019) or “*H. undulata*” (Appleyard 2020). Sub-clade *IV*_2_ specific to the Red Sea and adjacent localities was separated from sub-clade *IV*_1_ by 1.8% nucleotide distance at the *CO1* locus. The mean nucleotide distance was 0.3% within sub-clade *IV*_1_ and 0.4% within sub-clade *IV*_2_.

**Fig. 3.**
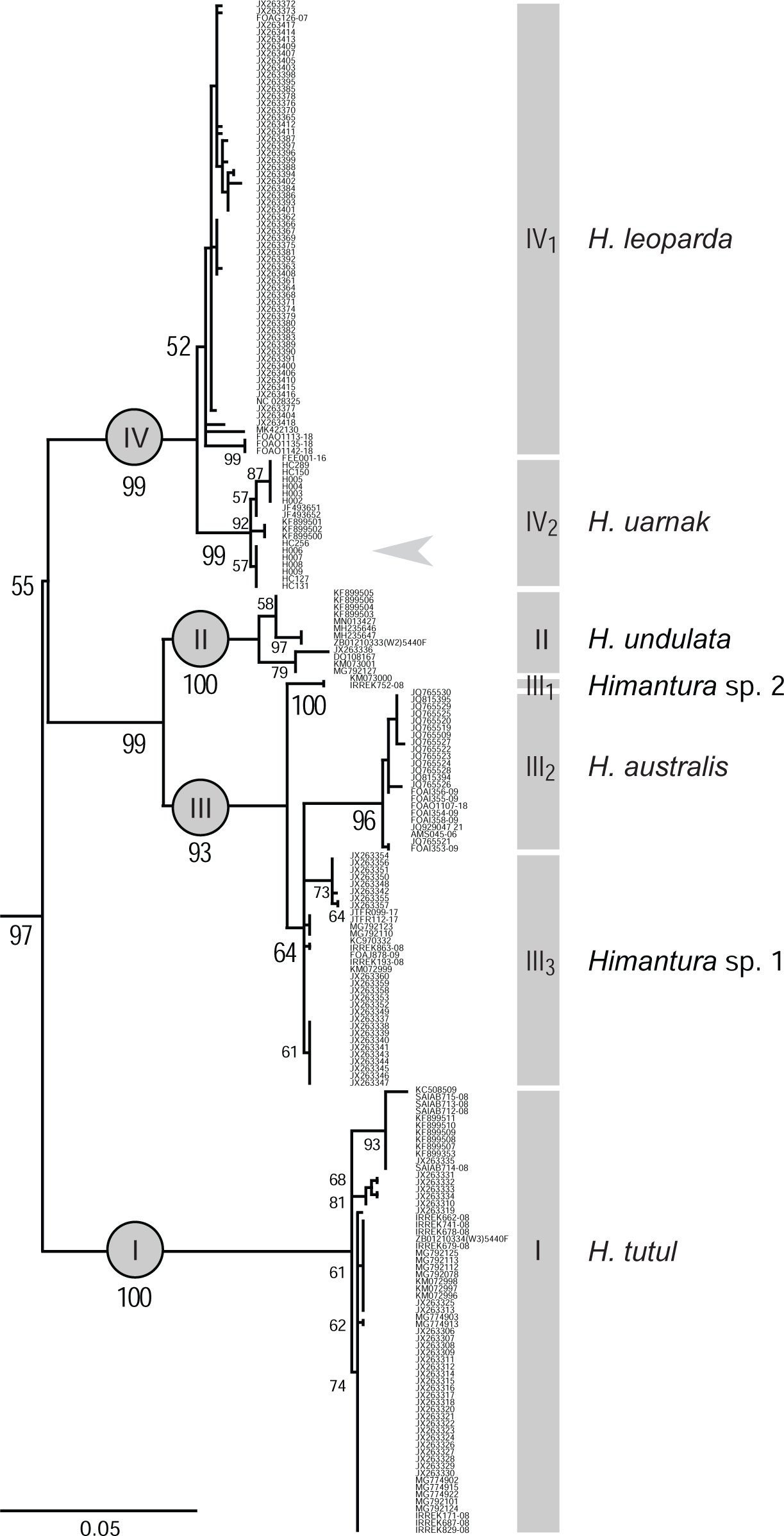
Reticulate whipray *Himantura* spp. species complex. Maximum-likelihood tree of *CO1* gene haplotypes based on the HKY+G+I model (MEGA X; Kumar et al. 2018). Tree with highest log likelihood (−16139.6) is shown. Tree is drawn to scale, with branch lengths measured in the number of substitutions per site. Haplotype sequences are listed using their GenBank or BOLD accession number. Scores at a node are percentages of trees in which the associated taxa clustered together (from 500 iterations of bootstrap resampling; Felsenstein 1985). The four main clades are designated by roman numbers *I-IV* following Arlyza et al. (2013); numbers in subscript indicate a sub-clade or an haplogroup within a clade. Arrow indicates placement of *H. uarnak* neotype

## Discussion

The taxonomic value of mitochondrial DNA sequences has been demonstrated in morphologically intractable species complexes in Elasmobranchs such as those of the long-tailed butterfly ray *Gymnura poecilura* (Naylor et al. 2012; Muktha et al. 2018), the whitespotted whipray *Maculabatis gerrardi* (Ward et al. 2008; Naylor et al. 2012) or the blue-spotted maskray *Neotrygon kuhlii* (Naylor et al. 2012; Puckridge et al. 2013; Borsa et al. 2018; Pavan-Kumar et al. 2018). The reticulate whipray *Himantura* spp. species complex is another example where morphological overlap between species has led to considerable confusion (Naylor et al. 2012; Borsa 2017), as confirmed by the plethora of names assigned to mitochondrial DNA haplotypes that cluster within the same clade or subclade (present survey).

In total, seven distinct lineages or haplogroups were identified in the phylogenetic tree of the reticulate whipray species complex (Fig. 3). Subclade *IV*_2_ included the *CO1* gene sequence of a specimen here chosen as *H. uarnak*’s neotype (see next section). As a consequence, sub-clade *IV*_2_ is here designated as P. Forsskål’s *H. uarnak*. The *H. uarnak* subclade was distinct from the subclade *IV*_1_ clustering all leopard whipray *H. leoparda* specimens from the Coral Triangle. *Himantura leoparda*, whose type locality is the Gulf of Carpentaria in the Coral Triangle region, thus is the sibling Pacific species of *H. uarnak*. The other lineages were identified as *H. australis* (sub-clade *III*_2_), *H. tutul* (clade *I*) and *H. undulata* (clade *II*) based on previous genetic work employing the *CO1* marker (Arlyza et al. 2013; Borsa et al. 2013; Borsa 2017; Appleyard 2020) and on geographic consistency with the type locality of *H. australis* (southern New Guinea; Last et al. 2016). Two sub-clades or haplogroups here labelled *Himantura* sp. 1 (haplogroup *III*_2_) and *Himantura* sp. 2 (subclade *III*_1_) remain to be described as new species or, possibly re-described as a resurrected species. Thus, the fixation of a neotype for *H. uarnak* clarifies the identity of over 200 specimens whose *CO1* gene sequences have been deposited in public databases and paves the way to the description or re-description of two other yet-unnamed species unnamed species in the genus *Himantura*.

Despite the brevity of its description, P. Forsskål’s *H. uarnak* has been universally treated as a single, valid taxon in the subsequent literature (Gmelin 1789; Rüppell 1835; Bleeker 1852; Duméril 1865; Last and Compagno 1999; Manjaji 2004). However, no type is known; and multiple lineages that qualify as distinct species have been assigned to *H. uarnak* (Naylor et al. 2012; Last et al. 2016; Borsa 2017) demonstrating taxonomic confusion. For clarifying past and future research dealing with the reticulate whipray species complex we elected to designate a neotype that unambiguously identifies *H. uarnak*.

### Neotype designation and re-description of *Himantura uarnak*

Genus *Himantura* Müller & Henle, 1837; species *H. uarnak* [Gmelin (ex-Forsskål), 1789], type species of the genus. The authorship of the species has been discussed extensively by Fricke (2008).

As argued above, designating a neotype for *Himantura uarnak* is necessary to clarify its taxonomic status. This act also helps secure the stability of the nomenclature of other species in the reticulate whipray species complex. Forsskål’s (1775) original description mentioned a whipray (“*cauda, quae apterygia*”) with spots all over (“*tota maculata*”). Reticulate whipray specimen KAUST-RSRC-H006 (male, 369 mm DW, collected on April 25, 2019 by AJM and CTW) has the foregoing attributes and it was captured off the Arabian shore of the central Red Sea ca. 80 km north of Jeddah, the assumed type locality (Fricke 2008). This specimen, which was allocated collection no. CAS-ICH 247241 in the fish collections of the California Academy of Sciences, San Francisco is here designated as *Himantura uarnak*’s neotype. This designation thus satisfies the conditions expressed in the International Code of Zoological Nomenclature (International Commission on Zoological Nomenclature 1999: Article no. 75).

Neotype locality is Thuwal, Saudi Arabia (22°18’40”N 39°05’26”E); habitat is shallow sand flat ∼50 cm deep adjacent to mangroves dominated by *Avicennia marina*.

Morphological description: disc rhomboidal, length 94.6% DW; snout with distinct apical lobe; anterior margins of disc convex, lateral apices narrowly rounded; posterior margin convex, free rear tip rounded. Pelvic fins moderately elongate, length 17.4% DW; width across base 14.5% DW. Tail whip-like, tapering gently toward sting, length 230.8% DW; see also Table 1, and Fig. 2a-d. Additional morphological measurements taken on neotype after it was fixed in formalin were the following (in mm): disc width 351, total length 1089, snout to pectoral-fin insertion 299, end of orbit to pectoral insertion 214, snout to maximum width 135, snout to origin of cloaca 284, cloaca origin to sting 129, pectoral insertion to sting origin 138, disc thickness 48, snout to preorbital (direct) 77, snout to preorbital (horizontal) 68, orbit diameter 16, eye diameter 19, spiracle length 14, orbit+spiracle length 36, interorbital width 51, inter-eye width 77, distance between spiracles 82, head length (direct) 165, preoral length (to lower jaw) 77, snout (prenasal) 58, nostril length 20, nasal curtain length 23, nasal curtain width 41, distance between nostrils 34, mouth width 38, distance between 1^st^ gill slits 74, distance between 5^th^ gill slits 51, width of 1^st^ gill slit 10, width of 3^rd^ gill slit 10, width of 5^th^ gill slit 7, tail width at axil of pelvic fins 21, tail width at base of sting 11, tail height at axil of pelvic fins 17, tail height base of caudal sting 8, sting length 79, cloaca length 2, greatest width across pelvic fins 88 (resting) or 102 (spread), clasper length 31 (post-cloaca) or 13 (from pelvic axil).

Genetic description: *CO1* gene sequences cluster into a distinct lineage in the phylogenetic tree of the genus *Himantura* (Fig. 3). Nucleotide sequence of partial *CO1* gene of neotype, comprised between homologous nucleotide sites no. 69 and no. 699 of the *CO1* gene in *H. leoparda* (GenBank accession no. NC_028325; Shen et al. 2016) is 5’-C G G T G C G T G A G C A C G G A T A G T G G G T A C T G G C C T T A G C C T G C T T A T T C G G A C A G A G C T A A G C C A A C C A G G C G C A T T A C T G G G T G A T G A T C A A A A A T A T A A T G T A A T T G T T A C C G C C C A T G C C T T C G T A A T A A T C T T T T T C A T G G T A A T A C C T A T T A T A A T T G G G G G C T T T G G T A A T T G A C T C G T C C C C C T A A T A A T C G G T G C **<u>T</u>** C C A G A T A T A G C C T T T C C T C G A A T A A A C A A C A T G A G T T T T T G A C T T C T T C C A C C A T C C T T T C T A C T A C T T T T G G C C T C T G C T G G A G T A G A G G C T G G C G C T G G A A C A G G C T G A A C A G T C T A T C C C C C A C T A G C T G G T A A T C T A G C A C A T G C A G G G G C T T C A G T A G A C T T A G C A A T C T T T T C C C T A C A C C T G G C C G G T G T A T C T T C T A T C **<u>C</u>** T **<u>A</u>** G C C T C T A T T A A T T T T A T C A C C A C A A T C A T T A A C A T A A A A C C A C C A G C A A T T T C G C A G T A T C A A A C A C C C C T C T T T G T C T G A T C A A T C C T T A T C A C A G C C G T A C T C C T C T T G T T A T C T C T T C C T G T C C T A G C A G C A G G T A T T A C **<u>G</u>** A T A C T T C T A A C A G A T C G T A A C C T C A A T A C A A C C T T C T T T G A T C C T G C A G G A G G A G G T G A C C C A A T T C T T T A T C A A -3’.

Four nucleotides here underlined characterize *H. uarnak*.

Geographic distribution: *H. uarnak* specimens whose identity was here ascertained from their partial *CO1* gene sequences (Supplementary Table S2) were previously reported as “*H. leoparda*” (Arlyza et al. 2013; Borsa et al. 2013; Yucel et al. 2017; Bineesh et al. 2018) or “*H. uarnak*” (Steinke et al. 2011). Conversely, the identity of specimens previously assigned to “*H. uarnak*” are now correctly assigned to *H. australis, H. tutul*, and two yet-unnamed *Himantura* spp. (see Supplementary Table S2). Based on these genetically validated records, the distribution of *H. uarnak* outside the Red Sea includes the eastern Mediterranean Sea (Yucel et al. 2017), Natal (Steinke et al. 2011), and the Arabian Sea (Bineesh et al. 2017).

## Acknowledgements

Foremost, thanks are due to M. Tietbohl for donating tissue samples, assisting in field collections, and providing laboratory assistance.. Additionally, Wwe are grateful to J. Spaet and members of KAUST’s Reef Ecology Lab for their support at various stages of this project; to D. Catania and L.A. Rocha for allocating a CAS collection number to the neotype of *H. uarnak*; to X. Chen for sharing reticulate whipray sequences; to M. Bennett-Smith for sharing excellent photographs; and to S. Bogorodsky, H. Debelius and R. Fricke for insightful discussions about the taxonomy and nomenclature of reticulate whiprays.

## Ethical approval

Reticulate whipray specimens were captured in accordance with KAUST’s institutional animal care and use committee (IACUC) under animal study proposal # 18IACUC14.

## Data availability

All data generated during this study are included in this published article or in the appended Supplementary Tables S1 and S2. Sequences produced through the present study have been deposited in GenBank (https://www.ncbi.nlm.nih.gov/nucleotide/) under accession nos. XX000000-XX000000. Photographs of all specimens are available from the authors upon request.

## Author contributions

AJM and CTW participated in the design of the study, collected and measured specimens, did the molecular analyses and participated in the writing. MLB contributed funding and laboratory facilities and supervised the study. PB designed the study, provided directions, compiled and analyzed the data and led the writing. All authors read, edited and approved the final version of the manuscript.

---

**Supplementary Table S1.** Sampling details for reticulate whipray *Himantura uarnak* specimens from the Red Sea including specimen no. KAUST-RSRC-H006 / CAS-ICH 247241 chosen as neotype (highlighted red)

| Specimen ID | Location | Date | Approximate depth | Collector | Method of capture | Sex | No. stings | Total length (cm) | Disk width (cm) | Snout-to-cloaca length (cm) |
| --- | --- | --- | --- | --- | --- | --- | --- | --- | --- | --- |
| H001 | Sirrayn south | 14 Feb. 2019 | 15 cm | AJM, CTW | 15-m seine net | F | 1 | 113.2 | 31.9 | 23.8 |
| H002 | Sirrayn south | 14 Feb. 2019 | 15 cm | AJM, CTW | 15-m seine net | F | 1 | 121.8 | 34.6 | 26.5 |
| H003 | Sirrayn south | 14 Feb. 2019 | 15 cm | AJM, CTW | 15-m seine net | M | na | 103.2 | 28.4 | 22.8 |
| H004 | Sirrayn south | 14 Feb. 2019 | 15 cm | AJM, CTW | 15-m seine net | F | 1 | 120.2 | 32.6 | 25.0 |
| H005 | Sirrayn south | 15 Feb. 2019 | 20 cm | AJM, CTW | 15-m seine net | M | 1 | 100.2 | 31.7 | 24.7 |
| H006 | KAUST north beach | 25 Apr. 2019 | 50 cm | AJM, CTW | 15-m seine net | M | 1 | 109.9 | 36.9 | 28.9 |
| H007 | Jeddah fish market | 29 Apr. 2019 | na | AJM, CTW | Chunk of pectoral fin purchased | na | na | na | na | na |
| H008 | KAUST north beach | 22 May 2019 | 20 cm | AJM, CTW | Handline baited with shrimp | M | 1 | 103.4 | 27.2 | 22.1 |
| H009 | KAUST harbor | Dec. 2016 | na | M. Tietbohl | 15-m seine net | F | 2 | na | 53.1 | na |
| HC127 | KAUST pier | 01 Feb. 2019 | 39 cm | AJM, CTW | 15-m seine net | F | na | 102.0 | 27.8 | 29.0 |
| HC131 | KAUST pier | 23 Mar. 2019 | 30 cm | AJM, CTW | Dip net | F | 2 | 113.4 | 31.0 | 24.6 |
| HC150 | KAUST north beach | 30 Apr. 2019 | 10 cm | AJM, CTW | 15-m seine net | F | 1 | 96.7 | 28.8 | 23.2 |
| HC256 | KAUST north beach | 25 Apr. 2019 | 80 cm | AJM, CTW | 15-m seine net | M | 1 | 197.0 | 74.8 | 54.8 |
| HC289 | KAUST pier | 01 Feb. 2019 | 74 cm | AJM, CTW | 15-m seine net | F | 2 | 184.0 | 58.0 | 48.0 |

**Supplementary Table S2.**
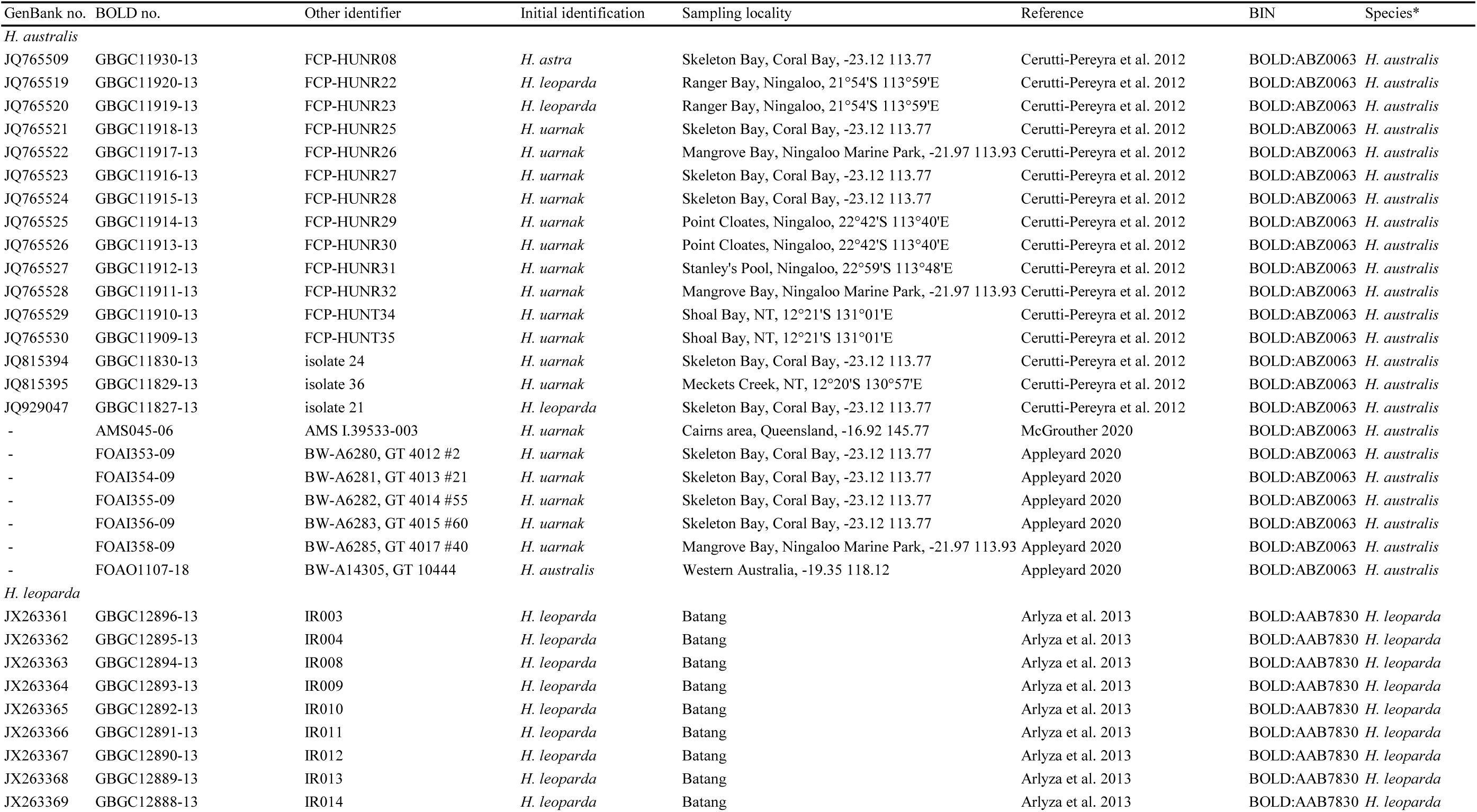
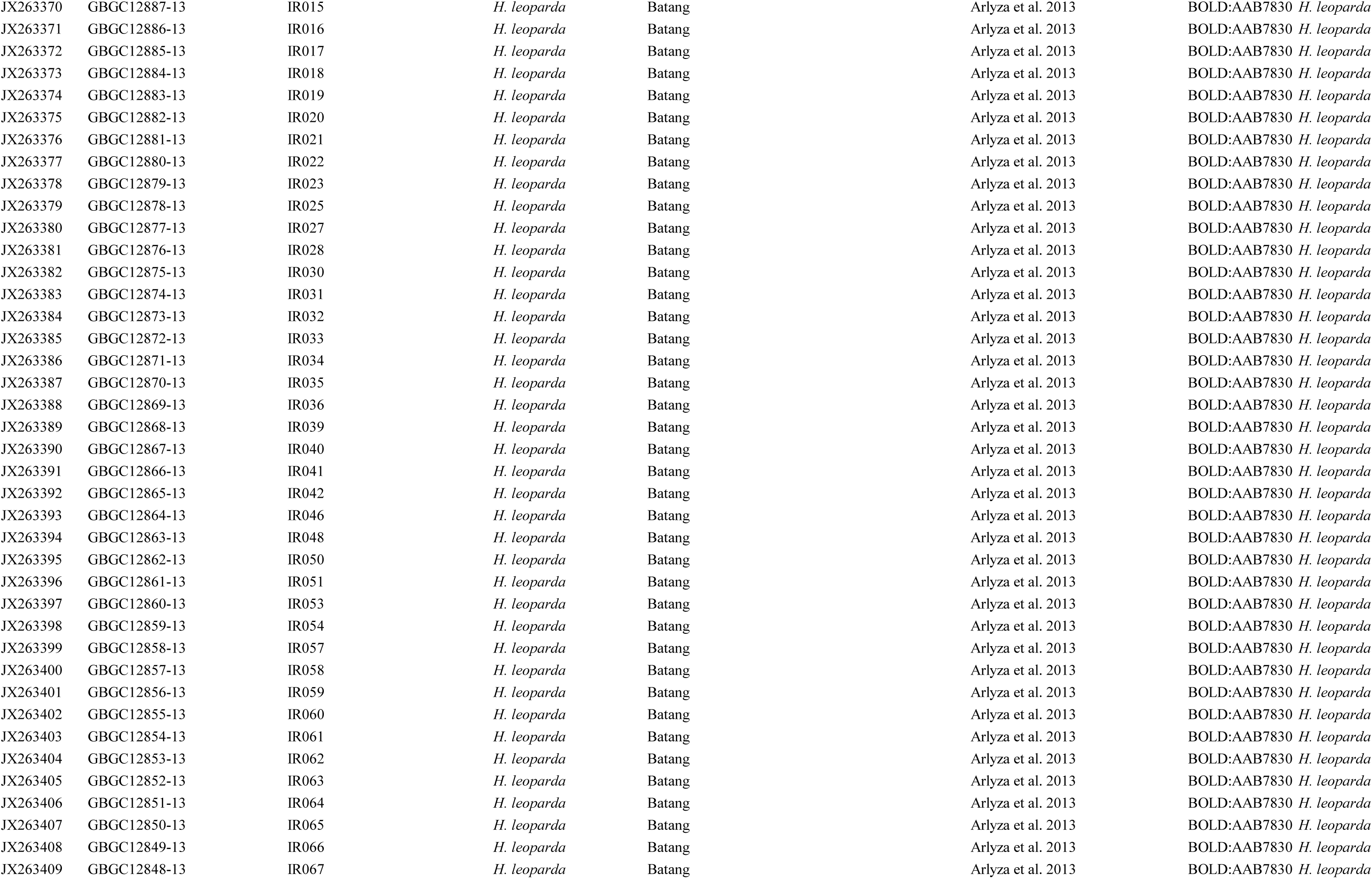
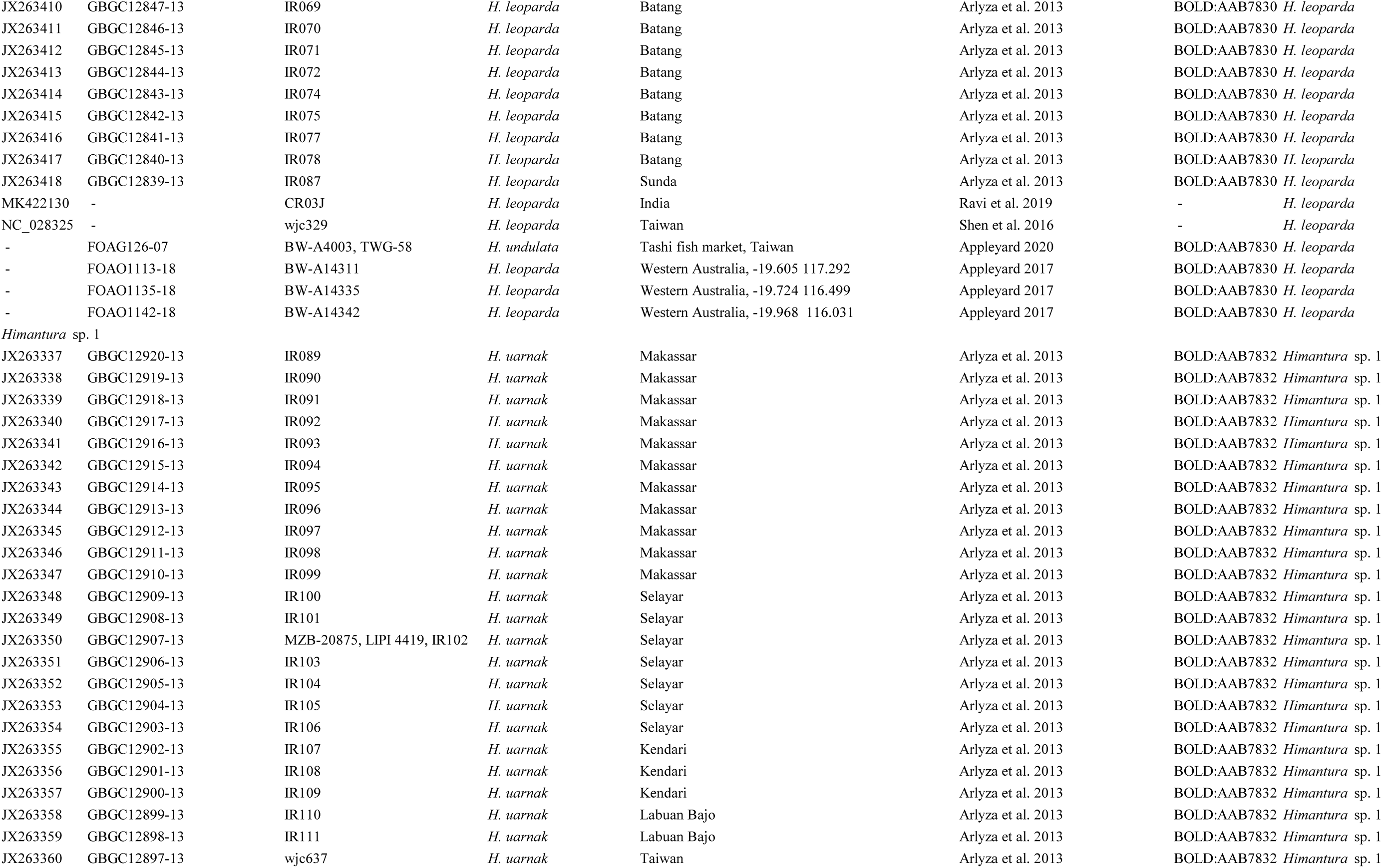
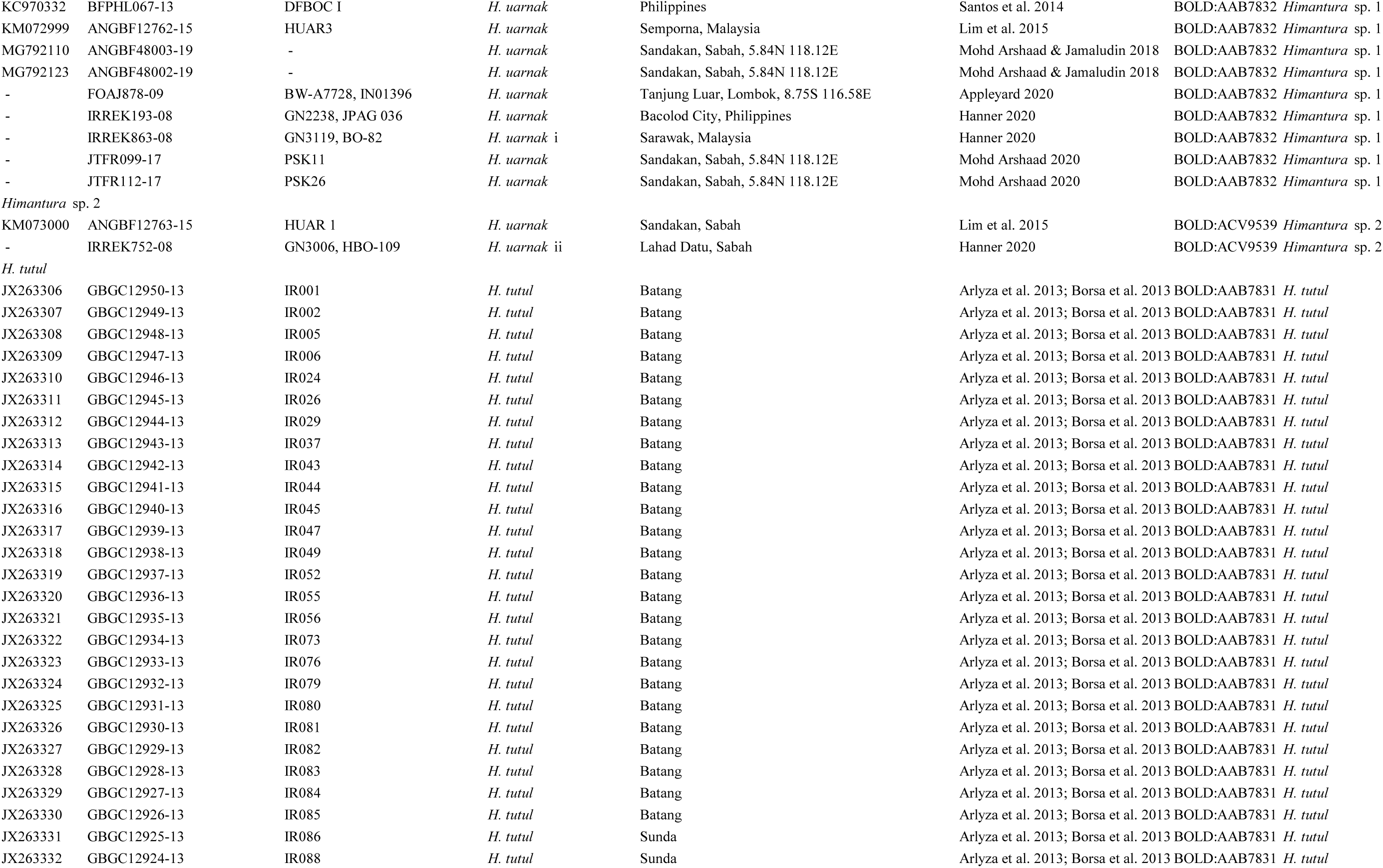
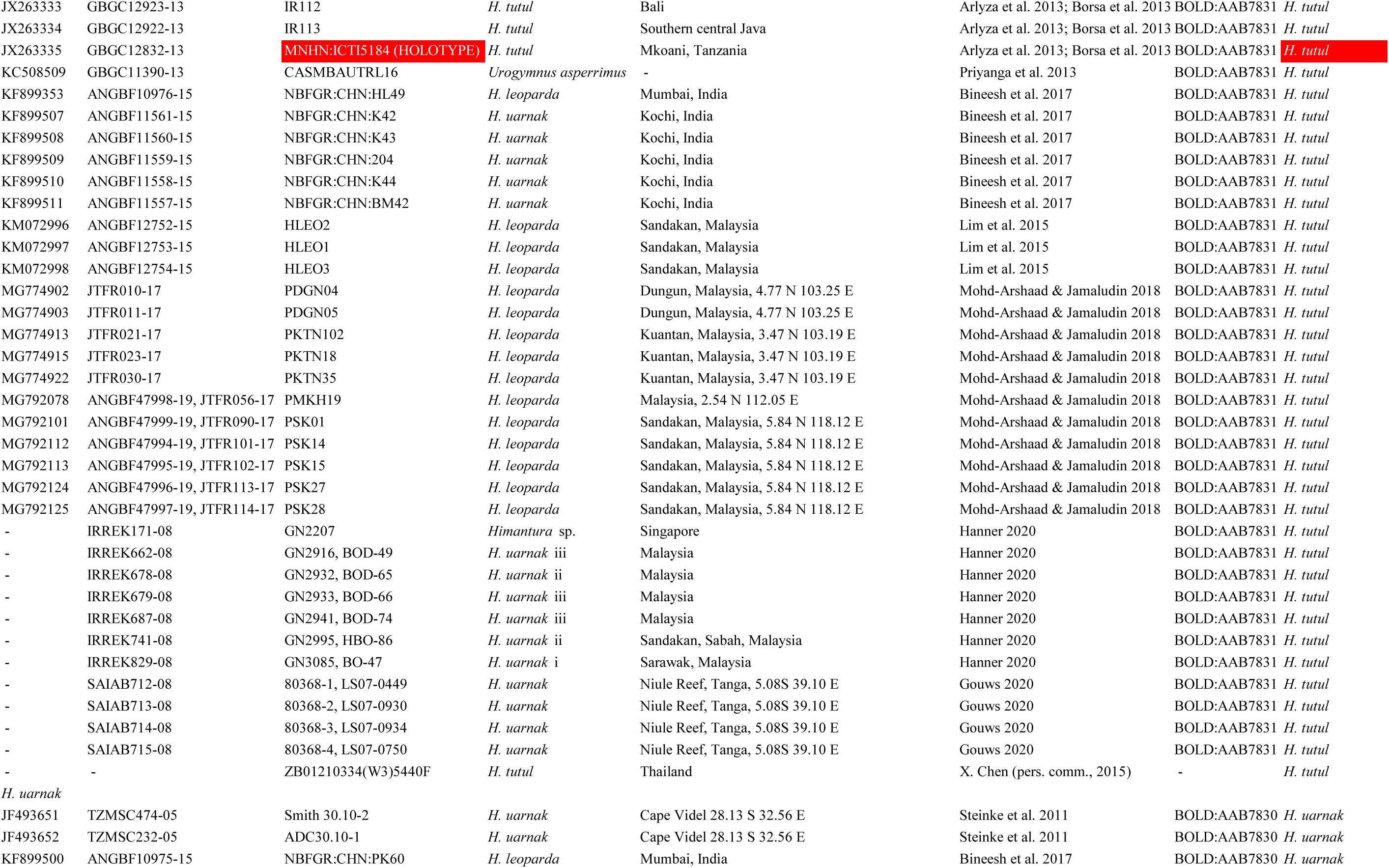
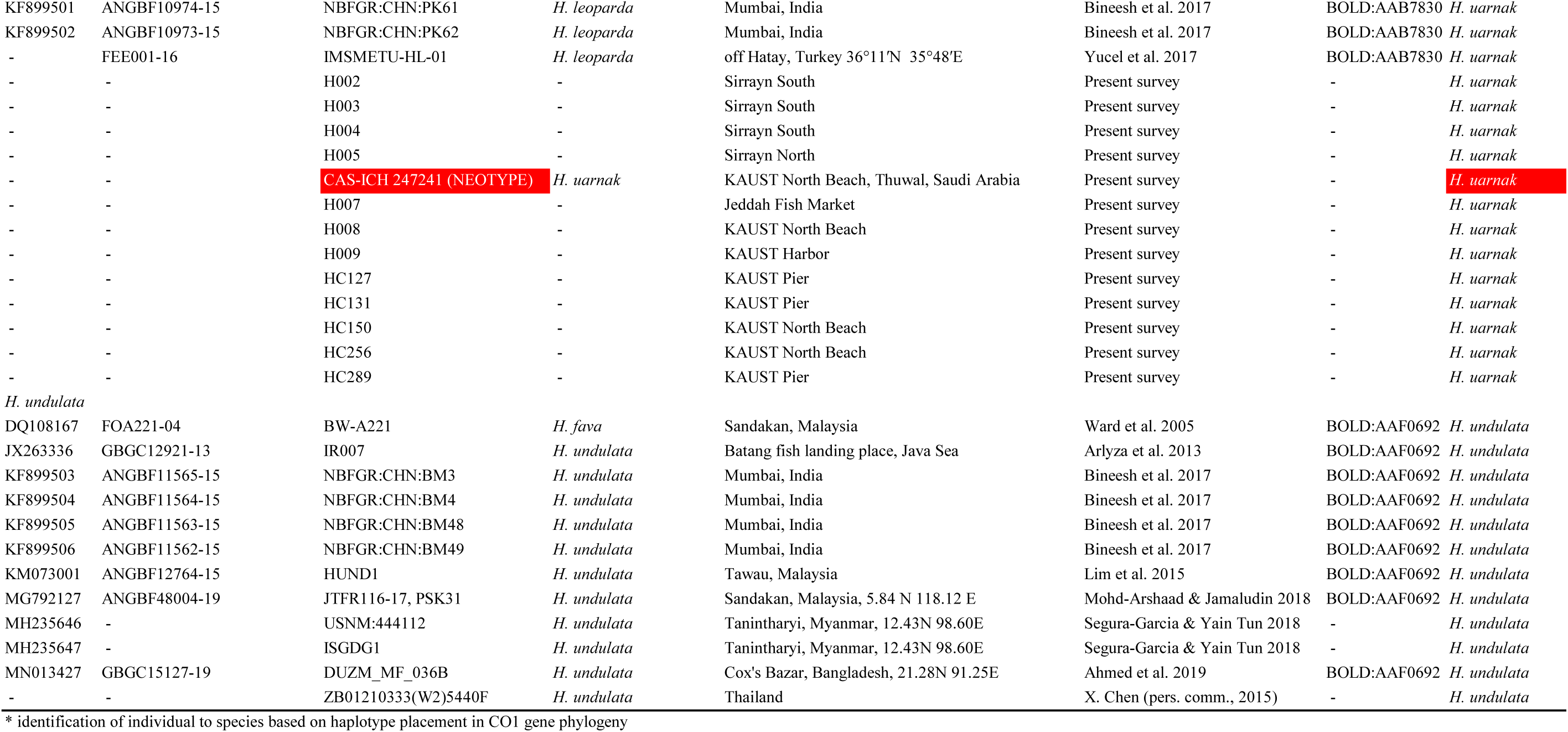
Reticulate whipray *Himantura uarnak* species complex. List of nucleotide sequences at the CO1 locus used for the present work, with GenBank and BOLD identification nos. and other details including sampling locations and references. Sequences were retrieved on 23 March 2020 from the GenBank and BOLD public databases (*N* = 201) or donated by X. Chen (Wenzhou Medical College, China) (*N* = 2) or produced through the present survey (*N* = 13). Sequences are arranged by species and presented in alphabetical order (*N* = 2) or produced through the present survey (*N* = 13). Sequences are arranged by species and presented in alphabetical order. Type material highlighted red

## References

Ahmed M.S., Zhilik A.A., Chowdhury N.Z., Ahmed S. (2019) DNA barcoding of marine fishes of Bangladesh. https://www.ncbi.nlm.nih.gov/nuccore/MN013427

Ali M, Saad A, Ben Amor MM, Capapé C (2010) First records of the honeycomb stingray, Himantura uarnak (Forskål, 1775), off the Syrian coast (eastern Mediterranean) (Chondrichthyes: Dasyatidae). Zool Middle East 49:104–106

Appleyard SA (2017) http://www.boldsystems.org/index.php/Public_RecordView?processid=FOAO1113-18, FOAO1135-18, FOAO1142-18

Appleyard SA (2020) http://www.boldsystems.org/index.php/Public_RecordView?processid=FOAG126-07, FOAJ878-09, FOAO1107-18

Arlyza IS, Shen K-N, Solihin DD, Soedharma D, Berrebi P, Borsa P (2013) Species boundaries in the Himantura uarnak species complex (Myliobatiformes: Dasyatidae). Mol Phyl Evol 66:429–435

Bineesh KK, Gopalakrishnan A, Akhilesh KV, Sajeela KA, Abdussamad EM, Pillai NGK, Basheer VS, Jena JK, Ward RD (2017) DNA barcoding reveals species composition of sharks and rays in the Indian commercial fishery. Mitoch DNA Pt A 28:458–472

Bleeker P (1852) Bijdrage tot de kennis der Plagiostomen van den Indischen archipel. Verhandel Batav Genoot Kunst Wetensch 24:1-92, pls 1-4

Borsa P (2017) Comments on “Annotated checklist of the living sharks, batoids and chimaeras (Chondrichthyes) of the world, with a focus on biogeographical diversity” (Weigmann, 2016). J Fish Biol 90:1170–1175

Borsa P, Durand J-D, Shen K-N, Arlyza IS, Solihin DD, Berrebi P (2013) Himantura tutul sp. nov. (Myliobatoidei: Dasyatidae), a new ocellated whipray from the tropical Indo-West Pacific, described from its cytochrome-oxidase I gene sequence. C R Biol 336:82–92

Cerutti-Pereyra F, Meekan MG, Wei N-WV, O’Shea O, Bradshaw CJA, Austin CM (2012) Identification of rays through DNA barcoding: an application for ecologists. PLoS One 7:e36479

Clark K, Karsch-Mizrachi I, Lipman DJ, Ostell J, Sayers EW (2016) GenBank. Nucl Acids Res 44:D67–D72

Duméril A (1865) Histoire naturelle des poissons ou ichthyologie générale. Tome premier Elasmobranches. Librairie encyclopédique du Roret, Paris, 720 p

Felsenstein J (1981) Evolutionary trees from DNA sequences: A maximum likelihood approach. J Mol Evol 17:368–376

Felsenstein J (1985) Confidence limits on phylogenies: an approach using the bootstrap. Evolution 39:783–791

Forsskål P (1775) Descriptiones animalium, avium, amphibiorum, piscium, insectorum, vermium; quae in itinere orientali observavit Petrus Forskål (post mortem auctoris edidit Carsten Niebuhr). Adjuncta est materia medica kahirina atque tabula maris Rubri geographica. Mölleri, Hauniae, xxxiv+164 p

Fricke R (2008) Authorship, availability and validity of fish names described by Peter (Pehr) Simon Forsskål and Johann Christian Fabricius in the ‘Descriptiones animalium’ by Carsten Niebuhr in 1775 (Pisces). Stuttg Beitr Naturk A Neue Ser 1:1–76

Gmelin JF (1789) Caroli a Linné […] Systema Naturae per regna tria naturae, secundum classes, ordines, genera, species; cum characteribus, differentiis, synonymis, locis. Editio decimo tertia, aucta, reformata, vol 1, pars III. G.E. Beer, Lipsiae, pp 1033–1516

Gouws G (2020) http://www.boldsystems.org/index.php/Public_RecordView?processid=SAIAB712-08, SAIAB713-08, SAIAB714-08, SAIAB715-08

Hall TA (1999) BIOEDIT: A user-friendly biological sequence alignement editor and analysis program for Windows 95/98/NT. Nucl Acids Symp Ser 41:95–98

Hanner R (2020) http://www.boldsystems.org/index.php/Public_RecordView?processid=IRREK171-08, IRREK193-08, IRREK662-08, IRREK678-08, IRREK679-08, IRREK687-08, IRREK741-08, IRREK752-08, IRREK829-08

Hasegawa M., Kishino H., Yano T. (1985) Dating the human-ape split by a molecular clock of mitochondrial DNA. J Mol Evol 22:160–174.

International Commission on Zoological Nomenclature (1999) International code of zoological nomenclature, 4th edn. International Trust for Zoological Nomenclature, London, 306 p

Kumar S, Stecher G, Li M, Knyaz C, Tamura K (2018) MEGA X: Molecular evolutionary genetics analysis across computing platforms. Mol Biol Evol 35:1547–1549.

Last PR, Compagno LJV (1999) Dasyatidae - Stingrays. In Carpenter KE, Niem VH (eds) FAO species identification guide for fishery purposes. The living marine resources of the Western Central Pacific. Volume 3. Batoid fishes, chimaeras and bony fishes part 1 (Elopidae to Linophrynidae). FAO, Rome, 1479–1505.

Last PR, White WT, Naylor G (2016) Three new stingrays (Myliobatiformes: Dasyatidae) from the Indo-West Pacific. Zootaxa 4147:377–402

Lim KC, Lim PE, Chong VC, Loh KH (2015) Molecular and morphological analyses reveal phylogenetic relationships of stingrays focusing on the family Dasyatidae (Myliobatiformes). PLoS One 10:e0120518

Manjaji BM (2004) Taxonomy and phylogenetic systematics of the Indo-Pacific whip-tailed stingray genus Himantura Müller and Henle 1837 (Chondrichthyes: Myliobatiformes: Dasyatidae). PhD Thesis, University of Tasmania, xxii+607 p

Manjaji-Matsumoto BM, Last PR (2008) Himantura leoparda sp. nov., a new whipray (Myliobatoidei: Dasyatidae) from the Indo-Pacific. CSIRO Mar Atmosph Res Pap 22:292–301

Manjaji-Matsumoto BM, White WT, Gutteridge AN (2016) Himantura uarnak. IUCN Red List Threat Sp 2016:e.T161692A68629130

McGrouther M.A. (2020) http://www.boldsystems.org/index.php/Public_RecordView?processid=AMS045-06

Mohd Arshaad W (2020) http://www.boldsystems.org/index.php/Public_RecordView?processid=JTFR099-17, JTFR112-17

Mohd Arshaad W, Jamaludin N-A (2018) DNA barcoding of rays in Malaysia. https://www.ncbi.nlm.nih.gov/nuccore/MG774902, MG774903, MG774913, MG774915, MG774922, MG792078, MG792101, MG792110, MG792112, MG792113, MG792123, MG792124, MG792125

Muktha M, Akhilesh KV, Sandhya S, Jasmin F, Jishnudev MA, Kizhakudan SJ (2018) Re-description of the longtail butterfly ray, Gymnura poecilura (Shaw, 1804) (Gymnuridae: Myliobatiformes) from Bay of Bengal with a neotype designation. Mar Biodiv 48:1085–1096

Naylor GJP, Caira JN, Jensen K, Rosana KAM, White WT, Last PR (2012) A DNA sequence-based approach to the identi?cation of shark and ray species and its implications for global elasmobranch diversity and parasitology. Bull Am Mus Nat Hist 367:1–262

Pavan-Kumar A, Kumar R, Pitale P, Shen K-N, Borsa P (2018) Neotrygon indica sp. nov., the Indian-Ocean blue spotted maskray (Myliobatoidei, Dasyatidae). C R Biol 341:120–130

Priyanga K, Thangaraj M, Singh R (2013) Molecular identification of Urogymnus asperrimus (family: Dasyatidae) using cytochrome oxidase I (COI) as a molecular marker.https://www.ncbi.nlm.nih.gov/nuccore/KC508509

Puckridge M, Last PR, White WT, Andreakis N (2013) Phylogeography of the Indo-West PAcific maskrays (Dasyatidae, Neotrygon): A complex example of chondrichthyan radiation in the Cenozoic. Ecol Evol 3:217–232

Randall JE (1983) Red Sea reef fishes. IMMEL Publishing, London, 192 p

Randall JE (1995) Coastal fishes of Oman. Crawford, Bathurst, xii+439 p

Ratnasingham S, Hebert PD (2007) BOLD: the barcode of life data system (http://www.barcodinglife.org). Mol Ecol Notes 7:p55–364

Ravi RK, Venu S, Bineesh KK, Akhilesh KV, Sajeela KA, Basheer VS, Gopalakrishnan A (2019) https://www.ncbi.nlm.nih.gov/nuccore/MK422130

Rüppell E (1835) Neue Wirbelthiere zu der Fauna von Abyssinien gehörig. Fische des rothen Meeres. Siegmund Schmerber, Frankfurt am Mein, ii+148 p, 33 pl

Santos MD, Asis AMM, Lacsamana JKM (2014) Illegal trade of regulated and protected aquatic species in the Philippines detected by DNA barcoding. https://www.ncbi.nlm.nih.gov/nuccore/KC970332

Segura-Garcia I, Yain Tun T (2018) Unscrambling Myanmar marine fishery biodiversity using DNA barcoding. https://www.ncbi.nlm.nih.gov/nuccore/MH235646, MH235647

Shen K-N, Chang C-W, Tsai S-Y, Wu S-C, Lin Z-H, Chan Y-F, Chen C-H, Hsiao C-D, Borsa P (2016) Next generation sequencing yields the complete mitogenomes of leopard whipray (Himantura leoparda) and blue-spotted maskray (Neotrygon kuhlii) (Chondrichthyes: Dasyatidae). Mitoch DNA Pt A 27:2613–2614

Steinke D, Zemlak TS, Connell AD, Heemstra PC, Hebert PDN (2011) https://www.ncbi.nlm.nih.gov/nuccore/JF493651, JF493652

Ward RD, Holmes BH, White WT, Last PR (2008) DNA barcoding Australasian chondrichthyans: results and potential uses in conservation. Mar Freshw Res 59:57–71

Ward RD, Zemlak TS, Innes BH, Last PR, Hebert PDN (2005) DNA barcoding Australia’s fish species. Phil Trans Roy Soc Lond B 360:1847–1857

Yucel N, Sakalli A, Karahan A (2017) First record of the honeycomb stingray Himantura leoparda (Manjaji-Matsumoto & Last, 2008) (Myliobatoidei: Dasyatidae) in the Mediterranean Sea, confirmed by DNA barcoding. J Appl Ichthyol 33:530–532

